# Object selection by automatic spreading of top-down attentional signals in V1

**DOI:** 10.1101/2020.02.24.962761

**Authors:** Matthias Ekman, Pieter R. Roelfsema, Floris P. de Lange

**Affiliations:** Radboud University Nijmegen, Donders Institute for Brain, Cognition and Behaviour, Nijmegen, Netherlands; Department of Vision and Cognition, Netherlands Institute for Neuroscience, an Institute of the Royal Netherlands Academy of Arts and Sciences, The Netherlands; Department of Integrative Neurophysiology, Center for Neurogenomics and Cognitive Research, VU University, Amsterdam, The Netherlands; Department of Psychiatry, Academic Medical Center, Amsterdam, The Netherlands

## Abstract

What is selected when attention is directed to a specific location of the visual field? Theories of object-based attention have suggested that when spatial attention is directed to part of an object, attention does not simply enhance the attended location but automatically spreads to enhance all locations that comprise the object. Here, we tested this hypothesis by reconstructing the distribution of attention from population neuronal activity patterns in V1 using functional magnetic resonance imaging (fMRI) and population-based receptive field mapping. We find that attention spreads from a spatially cued location to the underlying object – and enhances all spatial locations that comprise the object. Importantly, this spreading was also evident when the object was not task-relevant. These data suggest that attentional selection automatically operates at an object level, facilitating the reconstruction of coherent objects from fragmented representations in early visual cortex.

## Introduction

Imagine reaching for the coffee mug on your desk. Being able to grasp it properly requires precise information about which visual elements, like the mug’s handle, are part of the object and dissociate it from the background. In the visual system, object-based attention groups elements of spatially extended features, like the coffee mug, into one coherent object representation (Duncan, 1984; Egly et al., 1994). A likely grouping mechanism is the spreading of attention from a spatial location on the object that successively activates all parts of the object and thereby segregates it from the background and other objects (Roelfsema, 2006).

Recent studies showed that perceptual grouping according to Gestalt criteria occurs as early as in the primary visual cortex (V1) (Wannig et al., 2011; Pooresmaeili and Roelfsema, 2014). These previous studies, using multiunit recordings in non-human primates, focused on a limited number of receptive field locations, only covering parts of the attended and unattended objects (Wannig et al., 2011; Pooresmaeili and Roelfsema, 2014). It is therefore unclear whether V1 activity patterns carry a precise representations of the entire object, as postulated by theories of object-based attention, or rather represent only strategic object locations, while shape-selective neurons in higher visual areas represent the object with high fidelity (Moran and Desimone, 1985; Murray et al., 2002; Deco and Rolls, 2004). Elucidating this requires to investigate attentional spreading across the entire visual field representation within V1, which is typically not possible in multiunit recordings.

Here, we took advantage of the high spatial resolution of fMRI and recent advances in population based receptive field mapping to reconstruct the entire visual scene — including multiple objects (cued and non-cued) and the scene background, from V1 activity patterns. Stimulus reconstructions allowed us to directly test, with high spatial fidelity, whether attention spread from a cued location to the foreground object, but not the background object, as predicted by theories of object-based attention (see **Figure 1a** for an illustration).

**Figure 1.**
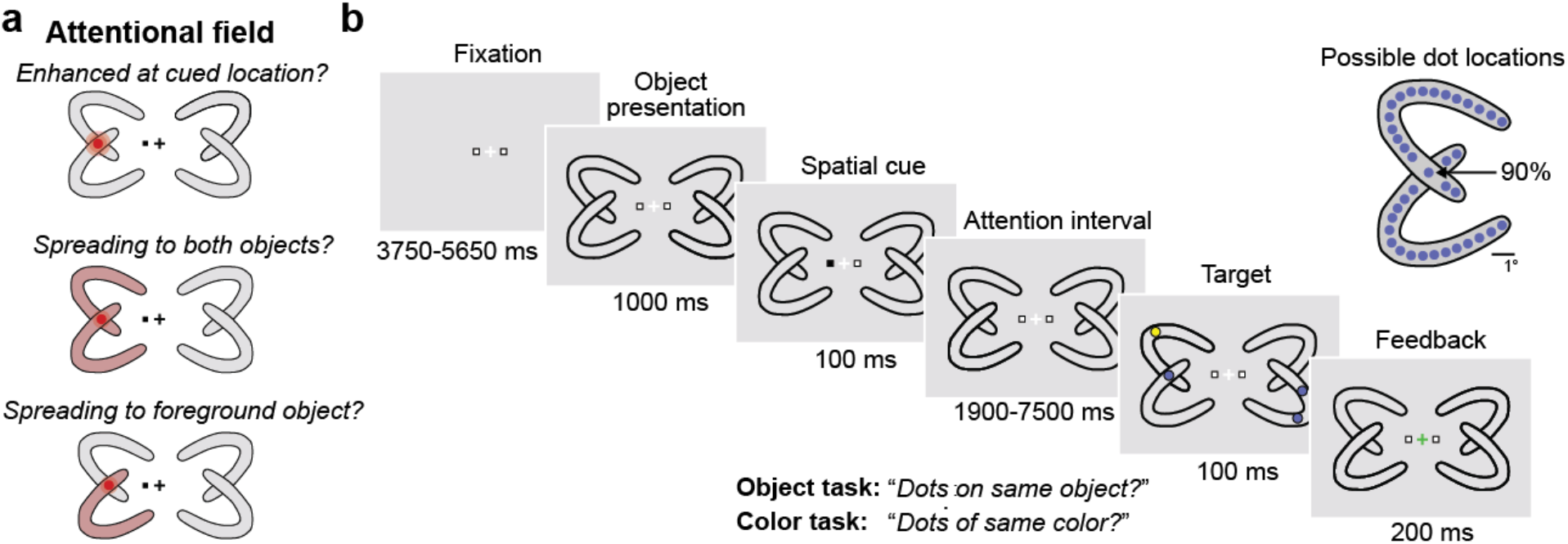
Experimental paradigm and hypothetical distributions of visual attention. (**a**) Hypothetical modulation of the attentional field. In this stimulus example the left hemifield is cued, causing changes in the distribution of attention at the cued location, at the intersection of the two objects. The spatial cue summons spatial attention to the precise position of the cued location (*top*). Attention spreads from the cued location to both objects (*middle*). Spread of the attention according to Gestalt criteria (*bottom*). (**b**) Trial schematic. Each trial begun with the presentation of four horseshoes followed by a spatial cue that indicated to attend either to the left or right hemifield. After a variable cue-to-target interval, four colored dots were presented on the objects: two relevant dots on the cued side and two irrelevant dots on the uncued side. The inset illustrates all possible dot locations. Based on the task instructions, participants had to either respond whether the dots on the cued side had the same color (*color task*), or whether they were located on the same object (*object task*). In 90% of all trials, one of the relevant dots appeared at the intersection of the two objects, and the second dot was equally likely to be on the same or different object. At the end of the trial, after a response window (1000 ms), feedback was presented in form of a green (correct response) or red (incorrect response) fixation cross. All fMRI analyses were restricted to the period before target onset. For illustration purposes, stimulus size and color were slightly adjusted. Whether a horseshoe was in the front or in the background was randomly chosen and counterbalanced across trials, for both the cued and uncued hemifield, respectively.

Furthermore, it remains an active debate whether object selection occurs automatically in the visual system (Müller and Kleinschmidt, 2003; He et al., 2004; Martínez et al., 2006; Yeari and Goldsmith, 2010; Wannig et al., 2011). For instance, perceptual grouping of the coffee mug might only occur when reaching for it, but not when typing on the nearby computer keyboard. Alternatively, perceptual object selection might require no voluntary control and occur automatically after attending a spatial location. We probed the automaticity of attentional spreading by varying the behavioral relevance of the objects in our paradigm (**Figure 1b**). We reasoned that if attentional spreading were also observed when the objects were task irrelevant, it would suggest that attentional spreading indeed occurs automatically.

In an exploratory, *second* fMRI experiment we focused on the temporal characteristics of attentional spreading using fMRI at higher sampling rate (TR=262 ms). We reasoned that if V1 activity successively spreads from the cued location along the curvature of the object, one would expect to see this process reflected in the temporal delay of BOLD responses along receptive fields on the object. Alternatively, if V1 activity spread from the cued location to the rest of the visual scene without following Gestalt criteria, one would expect a spatio-temporal BOLD profile that is more akin to a concentric wave instead of the object curvature.

To preview our results, group averaged stimulus reconstructions revealed that cued (but not uncued) objects showed enhanced V1 activity and were represented with high spatial acuity, supporting the idea that spatial attention spreads along the entire object. This neural enhancement was accompanied by a behavioral advantage for cued vs. uncued objects. Neural enhancement of the entire object was also present when the objects were not relevant, suggesting that attentional spreading occurs automatically. In sum, our findings indicate that attentional spreading constitutes a key mechanism for enhancing the detailed representation of objects in V1.

## Materials and Methods

### Data and code availability

All data and code used for stimulus presentation and analysis will be made available on the Donders Repository https://data.donders.ru.nl/ upon publication.

### Participants

A total of twenty-four healthy right-handed individuals (17 female, age 24 ± 3, mean ± SD) with normal or corrected-to-normal vision gave written informed consent to participate in this study, in accordance with the institutional guidelines of the local ethics committee (CMO region Arnhem- Nijmegen, The Netherlands). Seventeen subjects participated in experiment 1 and seven subjects participated in experiment 2.

### MRI acquisition parameters

Functional images for experiment 1 were acquired using a 3T Skyra MRI system (Siemens, Erlangen, Germany) with a T2*-weighted gradient-echo EPI sequence (TR/TE = 1800/30 ms, 26 transversal slices, voxel size 2×2×2 mm, 60° flip angle). Functional images for experiment 2 were acquired using a 3T Prisma MRI system (Siemens, Erlangen, Germany) with a multiband 3 sequence (TR/TE = 262/35.80 ms, 9 transversal slices, voxel size 2.4×2.4×2.4 mm, 38° flip angle, 20% slice gap). Anatomical images were acquired with a T1-weighted MP-RAGE sequence (TR/TE = 2300/3.03 ms, voxel size 1×1×1 mm, 8° flip angle).

### Visual stimuli and task

The objects consisted of two partially overlapping horseshoe-like objects in the left and right visual field, respectively. To the left and right of the fixation cross (0.4°), two white squares (0.05°) were shown throughout the trial. On each trial, one of the two squares changed from white to black for 100 ms cueing participants to attend either to the left or to the right visual field. The cue alternated between the right and left hemifield. Two colored dots (0.5°) appeared on the attended objects and also on the unattended objects (**Figure 1b**). During the *object-condition*, participants had to respond whether the two dots were on the same object, irrespective of the dot color. During the *color-condition*, participants had to indicate whether the dots were of the same color (yellow or blue), irrespective of whether the dots were on the same object. Participants were instructed to maintain fixation throughout the entire experiment.

A target dot was presented at 1 out of 19 different target locations per object. Notably, one of the dots on the attended side had an 88% chance (112 out of 128 trials) of occurring at the intersection of the horseshoe-objects (**Figure 1b**). This high probability of target appearance is expected to attract attention to that location (Shomstein and Yantis, 2004). The appearance of a target dot on any of the other 18 dot locations was equally likely. The object intersection belonged in 50% of all trials to the lower horseshoe, and in 50% to the upper horseshoe. The cued location did not predict whether the second dot would occur on the same or on the other object. The given stimulus parameters were the same for the cued and the uncued hemifield.

### Experimental design

The experiment was divided in four parts: 4 task runs, 1 localizer run, 4 retinotopic mapping runs and 1 anatomical scan. Each task run consisted of 64 trials and lasted for 13.4 min. At the beginning of each run, participants were instructed to perform the object or color task. The run order (i.e., two color tasks followed by 2 object tasks, or vice versa) was randomized across participants. Each trial started with a variable fixation period (average duration: 4700 ms, ranging from 3750-5650 ms), followed by presentation of the objects (1000 ms). A spatial cue (100 ms) indicated to attend to the left or right side and after a variable cue-target interval (average duration: 4700 ms, ranging from 1900-7500 ms), the target dots were shown for 100 ms. Participants were instructed to respond with the right-hand index and middle finger (index finger: ‘same’, middle finger: ‘different’). After a response window of 1000 ms feedback was given. Depending on the response, the fixation cross turned green (correct response), or red (incorrect response) for 200 ms. Each trial lasted on average 11.8 s. Additionally, six null-events (8.6 s), only showing the fixation cross, were presented at random positions per run, allowing the BOLD time-courses to return to baseline.

During the functional localizer, one of the four horseshoe objects was flashed for 500 ms on, 500 ms off, for 10.8 s per block. Each object block was presented four times in total and object presentations were separated by null events (8.6 s). During the localizer, participants saw a sequence of rapidly changing letters at fixation and had to report whenever target letters ‘X’ or ‘Y’ (target probability = 10%) appeared in a stream of non-target letters (‘A’, ‘T’, ‘N’, ‘U’, ‘V’, ‘Y’, ‘H’, ‘R’). Letters were presented for 400 ms each, separated by 400 ms intervals in which only the fixation point was presented.

### fMRI data preprocessing

All fMRI preprocessing was carried out using FSL (Smith et al., 2004). The first six volumes of each run were discarded to allow for stabilization of the magnetic field. All functional images were spatially realigned to the middle image of the first run to correct for head movement. The functional data were high-pass filtered (cutoff: 128 s) to remove low-frequency signal drifts. The functional volumes were aligned with the anatomical image using linear registration.

### Population receptive field measurements

After the main experiment, we presented moving bar stimuli, in order to map the population receptive fields (pRFs) of voxels in early visual cortex, as well as allow polar angle mapping to delineate the borders between retinotopic areas in visual cortex (Sereno et al., 1995; Engel et al., 1997). During these runs, bars containing full contrast flickering checkerboards (2 Hz) moved across the screen in a circular aperture with a diameter of 20°. The bars moved in eight different directions (four cardinal and four diagonal directions) in 20 steps of 1°, one step per TR (1.8 s). Four blank fixation screens (10.8 s) were inserted after each of the cardinally moving bars. Throughout each run (5.76 min), a colored fixation dot was presented in the center of the screen, changing color (red to green and green to red) at random time points. Participants’ task was to press a button whenever this color change occurred. Participants performed four identical runs of this task.

### pRF estimation

The data from the moving bar runs were used to estimate the population receptive field (pRF) of each voxel in the functional volumes using MrVista (http://white.stanford.edu/software/). In this analysis, a predicted BOLD signal is calculated from the known stimulus parameters and a model of the underlying neuronal population. The model of the neuronal population consisted of a two-dimensional Gaussian pRF, with parameters *x0*, *y0*, and *s0*, where *x0* and *y0* are the coordinates of the center of the receptive field, and *s0* indicates its size (standard deviation). All parameters were stimulus-referred, and their units were degrees of visual angle. These parameters were adjusted to obtain the best possible fit of the predicted to the actual BOLD signal. For details of this procedure, see (Dumoulin and Wandell, 2008). This method has been shown to produce pRF size estimates that agree well with electrophysiological receptive field measurements in monkey and human visual cortex. Once estimated, *x0 and y0* were converted to eccentricity and polar-angle measures. For the following analyses, only voxels with a model fit of R^2^ > 1% were considered. All analysis scripts for the pRF estimation are available here: https://github.com/mekman/pRF.

### Selection of V1 voxels

Area V1 was determined using the automatic cortical parcellation provided by Freesurfer 6.0 (Fischl, 2012) based on individual T1 images. With increasing receptive field size, voxels will respond to multiple dot locations. In order to prevent overlap in response profiles the V1 mask was further restricted to voxels with a pRF-size >= 3.5°.

### Attentional field reconstruction

We developed a multivariate pRF-based reconstruction method (**Figure 3**) that allows for more fine-grained reconstructions compared to the standard, summation-based pRF reconstruction (Thirion et al., 2006; Kok and De Lange, 2014; Ekman et al., 2017). First, a general linear model (GLM) was used to fit individual BOLD responses and obtain estimates of signal change per voxel. The GLM consisted of 4 regressors of interest (object presentation followed by attention interval: cue (left/right) and horseshoe (foreground/background)), target presentation (1 regressor), and 16 motion regressors. Note that both the ITI and the cue-target interval were temporally jittered, which aids the separate estimation of these events. Color and motion tasks were analyzed separately.

**Figure 2.**
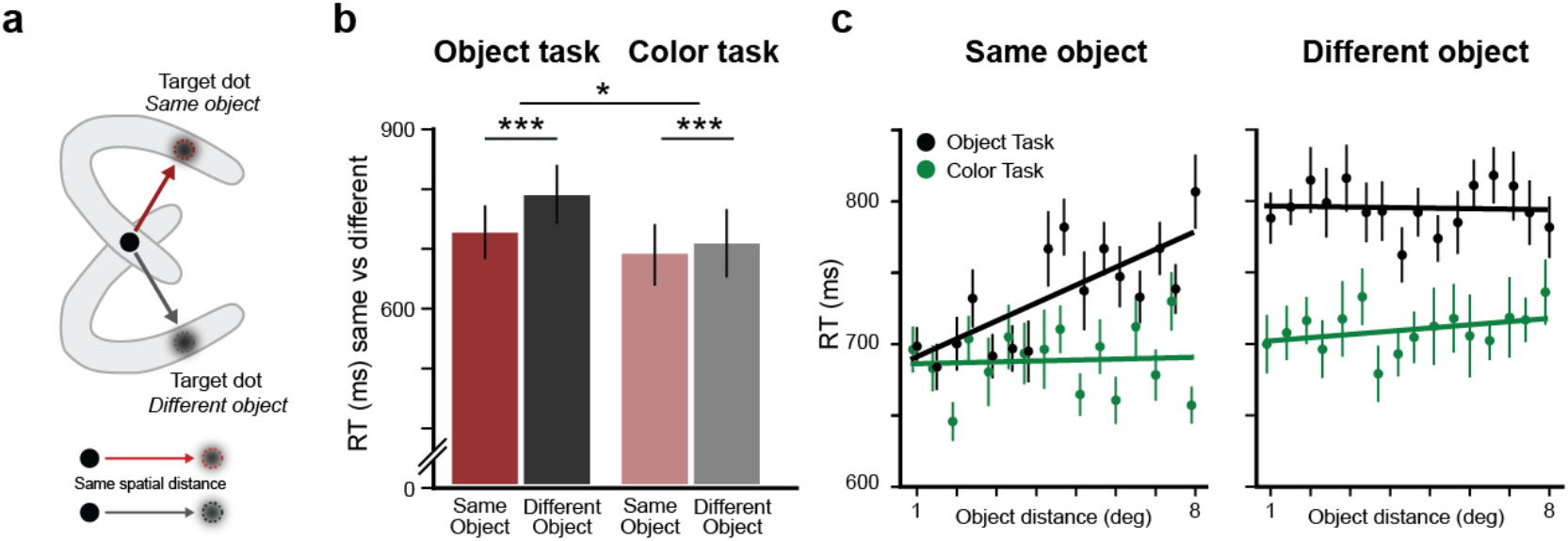
Reaction times in the two tasks. (**a**) Targets located on the same vs. different object have the same spatial distance from the cued location. (**b**) In the object task, RTs were longer if the targets were on different objects, as in previous studies (Jolicoeur et al., 1986). In the color task, a similar effect was found, even though the objects were irrelevant. (**c**) RTs as function of the location of the second target in the object task (*left*) and color task (*right*). Error bars denote s.e.m; \**P* < 0.05; \*\**P* < 0.01; \*\*\**P* < 0.001.

**Figure 3.**
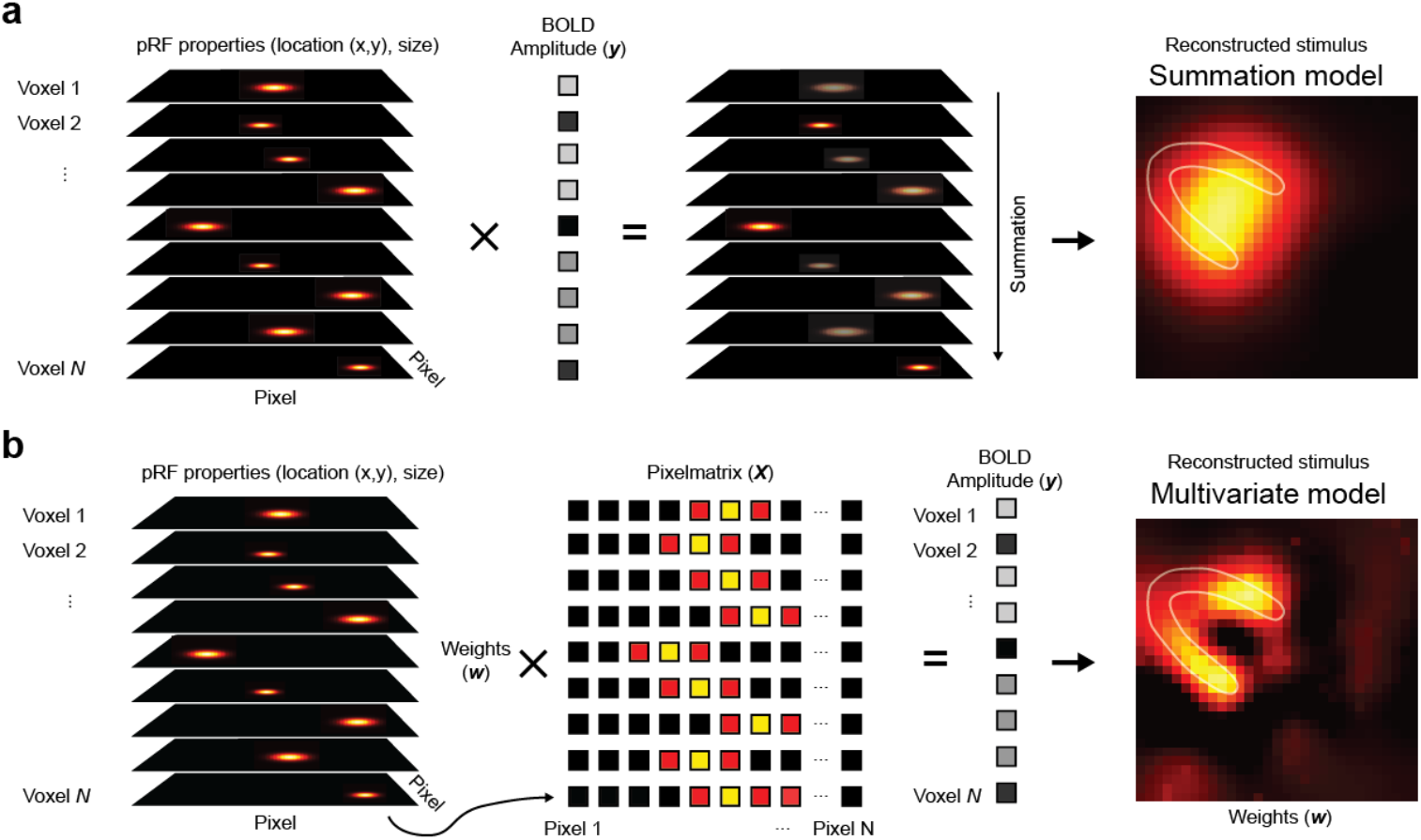
Illustration of the prf-based reconstruction analysis. (**a**) The standard model of prf-based reconstruction scales the prf properties for each voxel (x0, y0, s0) with the BOLD amplitude observed during each task condition. The reconstructed image is obtained by summing over the voxel dimension. (**a**) The multivariate prf-based reconstruction approach developed in this study follows three steps. First, the given receptive field properties for every voxel are represented in a 2D pixel matrix (*see Materials and Methods*). Second, the pixel matrix is then regressed against the measured BOLD amplitude, separately for each task condition. Third, plotting the estimated regression weights reveals the stimulus reconstruction. Stimulus reconstruction shown for the group averaged (N=17) left upper horseshoe from the localizer block. White contour lines illustrate the stimulus locations.

Second, every voxel is described as a 2D Gaussian with parameters *x0, y0*, and *s0* from the pRF estimation. In the standard pRF reconstruction (**Figure 3a**) the 2D Gaussians for each voxel, represented by a pixel × pixel image, are scaled based on the percent signal change, and consecutively summed over voxels to create one 2D representation of the reconstructed stimulus. One potential downside of this summation method is that a small number of reliably activated, or deactivated voxels might not show in the final reconstruction if they are surrounded by a larger number of deactivated, or activated voxels. In order to address this limitation we developed an alternative reconstruction approach, which allows us to reconstruct stimulus details by taken reliably activated voxels into account even if they are not present in a large quantity (**Figure 3b**). This is achieved by re-weighting the contribution of receptive-fields using a regression model. To this end, the 2D Gaussian describing each voxel (represented in this case by 21 × 21 image pixel) was flattened into a one-dimensional vector consisting of 21 × 21 = 441 pixels. An odd number of pixel was chosen so that each hemifield is covered by an equal number of pixel, i.e. 10 pixel for the left and 10 pixel for the right hemifield. Given these parameters, each pixel corresponds to roughly 0.76 degree of visual angle. For reference, the 2D surface area of a voxel with a receptive field size of 1° covers approximately 21 pixels, and a voxel with a receptive field size of 2° covers 89 pixels. Voxel with a receptive field < 0.4° were covered by only one single pixel. Given the median receptive field size of 1.44° for selected V1 voxel in this study, the 21 × 21 pixel grid ought to provide sufficient resolution for our reconstruction analysis. In fact, explorative analyses using data exclusively from the independent localizer revealed no advantage for the stimulus reconstruction when using a higher resolution, like a 31 × 31 pixel grid.

Let the matrix describing the voxels *v*, by pixel *p* be called *X*. The voxels’ signal change (in one particular task condition) is described in a vector *y*. Next, we performed Linear least squares regression to solve *X* for *y*. In order to deal with the multicolinearity of the receptive field data, we used ridge regression (Pedregosa et al., 2011) to estimate the weights *w*. The weights in the context of ridge regression are given by:

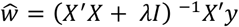

This formulation has a free ridge parameter λ, that was set to the number of selected voxels in the respective V1 mask. Initial exploration using data exclusively from the independent localizer revealed that determining the ridge parameter λ using an exhaustive grid search (values ranging from 0, corresponding to an ordinary least-squares regression, to 1000 in steps of 20) did not improve the reconstruction results visually. We therefore decided not to employ any grid search analysis.

Finally, the estimated weights (shape: 1 × 441) were reshaped into the original pixel shape of the 2D Gaussian (21 × 21 pixel in this case) and reflect the reconstructed image (one per regressor of interest; **Figure 4a**). The resulting 4 reconstructions per task condition (cue (left/right); horseshoe (foreground/background)) were combined into one reconstruction by rotating and flipping all reconstructions into the upper left-hand corner (**Figure 4b**) and we then computed the attentional modulation in percent, by comparing the attended and unattended hemisphere. The python code for the multivariate pRF reconstruction is available at https://github.com/mekman/recon.

**Figure 4.**
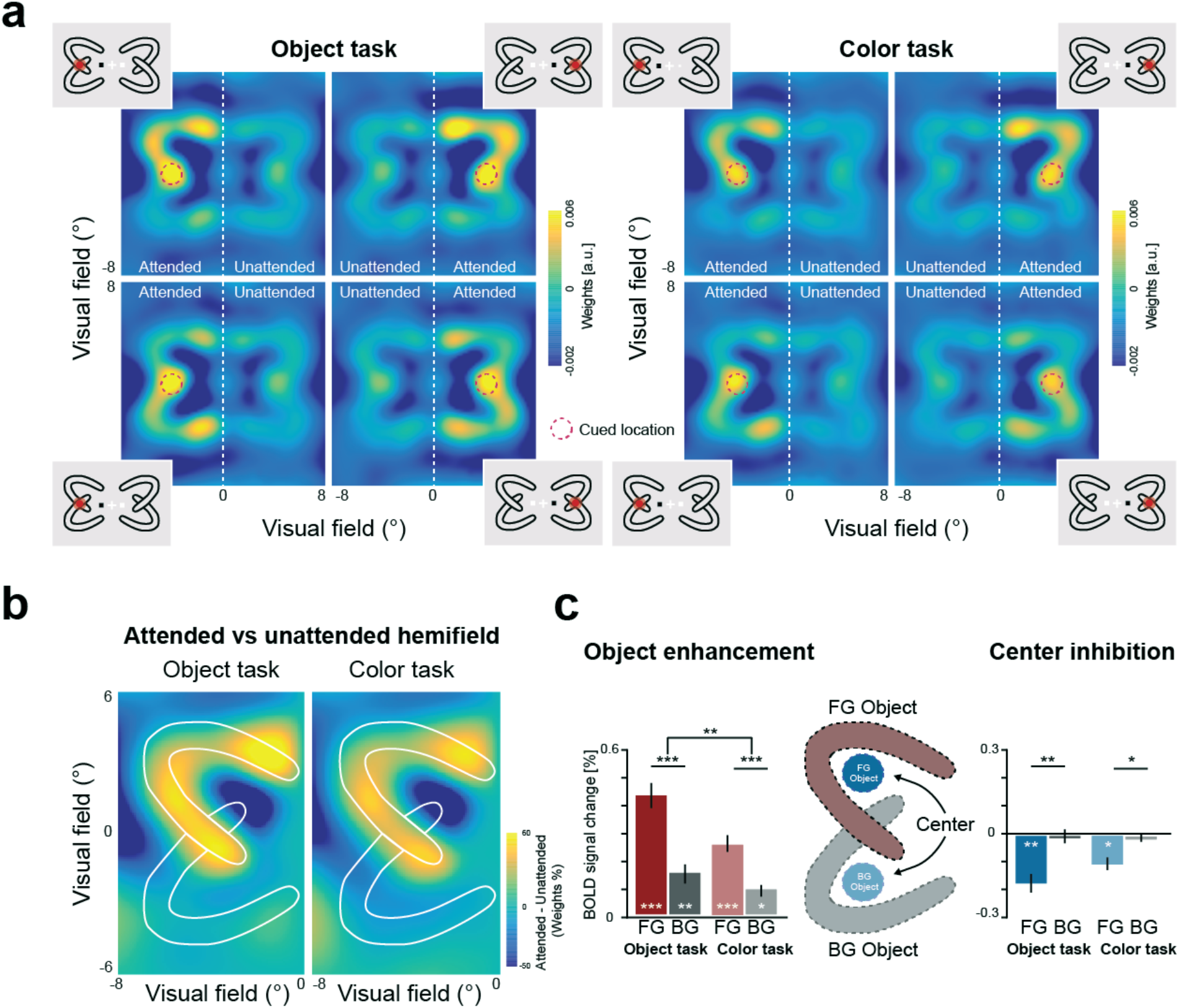
Attention spreads across the foreground object on the cued side. (**a**) Receptive field-based stimulus reconstruction from BOLD activity in V1. Group-averaged stimulus reconstructions (N=17 participants) in response to the spatial cue for all four stimulus configurations, separately for the object task (*left*) and color task (*right*). The red circle highlights the high probability target location. (**b**) The four stimulus configurations were combined into one image by flipping all cued foreground objects into the upper left corner. The attentional effect was calculated as the BOLD activity difference between the attended and unattended hemifield. White contour lines indicate the stimulus location. Attention spread from the cued location over the underlying object according to Gestalt criteria. The effect was found when the object was relevant for the task (i.e., object task; *left*) and when the object was not relevant to the task (i.e., color task; *right*). (**c**) Bar plots show the averaged BOLD activity for voxels with a receptive field on the cued object, compared to the non-cued object (both in the cued hemifield), separately for the object task and color task (*left*). The BOLD activity was higher on the foreground (FG) object than on the background (BG) object. Illustration of receptive field locations that were selected for this analysis (foreground (FG) object, background (BG) object, foreground (FG) object center, and background (BG) object center; *middle*). The bar plots show the averaged BOLD activity for voxels with a receptive field on the FG object center, compared to the BG object center, separately for the object and color task (*right*). Error bars denote s.e.m; \**P* < 0.05; \*\**P* < 0.01; \*\*\**P* < 0.001.

Voxels on the foreground and background object were selected based on their voxel receptive fields and mean BOLD percent signal change estimates were obtained by averaging across voxels, separately for the object- and color task. Further, voxels with receptive fields at the center of the horseshoe-like objects were selected, separately for the object- and color task to test for possible center inhibition effects. Participants’ mean BOLD estimates were compared using ANOVA.

### HRF duration

In order to test for differences in the width (i.e., dispersion) of BOLD time-courses between the color and the motion task, we additionally employed finite impulse response (FIR) modeling (Ollinger et al., 2001) as implemented in FSL FEAT. Resulting hemodynamic response function (HRF) estimates were averaged across voxels with a receptive field on the foreground object, separately for the color and motion task. Subsequently, a standard single gamma function with three parameters, amplitude, peak-time and FWHM (i.e., dispersion) was used to fit individual HRF responses (Glover, 1999). Finally, HRF dispersion parameters of the color and motion task were compared using paired sample t-test.

### Temporal spreading

fMRI response amplitude and peak latency were computed by fitting a conventional single gamma HRF function (Boynton et al., 1996) to individual voxel time-courses for pRF locations ‘P1’ and ‘P2’ (see **Figure 6**) for each participant and task condition. This was done using the function *curve_fit* as implemented in SciPy 0.18 (based on the Trust Region Reflective algorithm for a constrained least-squares fit). The objective function was the sum of squared errors between the predicted and observed response. We allowed the baseline, amplitude and peak delay to vary as free parameters. The peak delay was constrained to a range between 3-11.5 s peak latency in order to prevent pathological fits. The resulting peak latency for P1 and P2 was compared with a two-sided *t*-test separately for the same- and different object. Due to the low number of participants in experiment 2 we additionally performed a Bayesian *t*-test using JASP (JASP Team, 2019).

**Figure 5.**
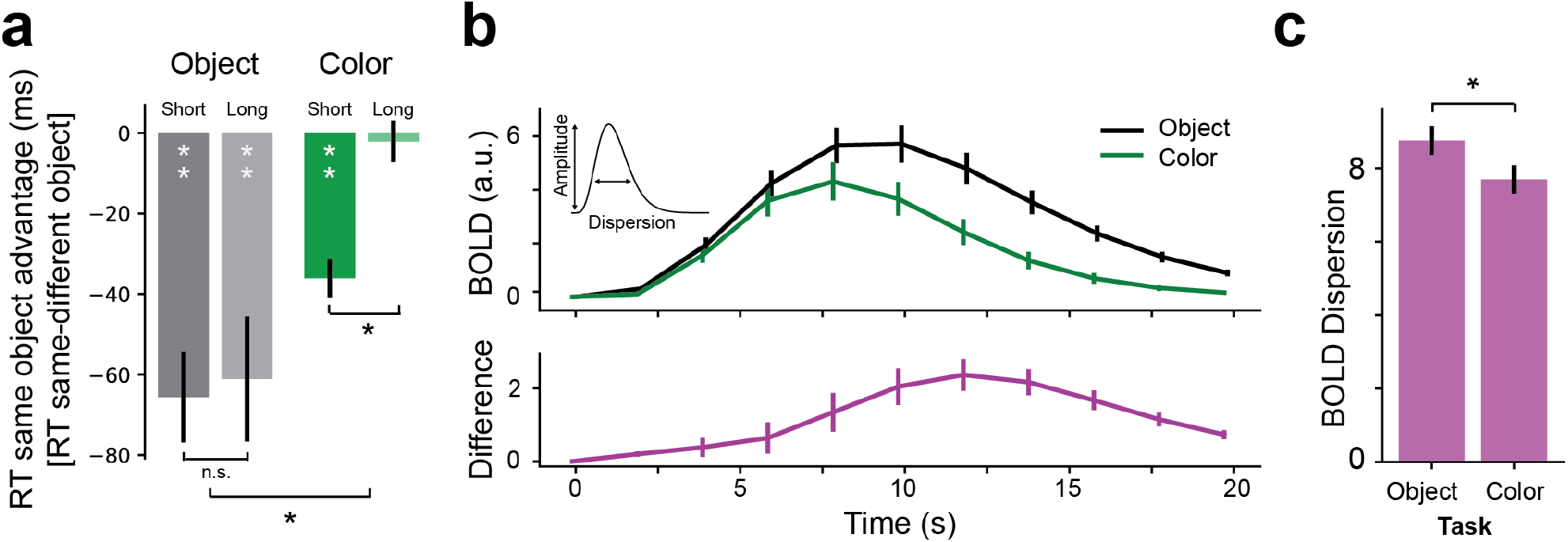
Task context modulates temporal dynamics of attentional object selection. (**a**) RT on trials with targets on the same object minus RT with the targets on different objects for shorter and longer cue-target delays. In the object task RTs are always shorter when both cues appear on the same object. In contrast, in the color task, the RT advantage is present for shorter – but not for longer – cue-target delays, suggesting that the attention selection of the foreground object is not maintained throughout the delay. (**b**) Group-averaged deconvolved BOLD time-courses (N=17) for V1 voxel with receptive fields along the cued object (*top*) during the object task (*black*) and color task (*green*). Difference profile of the BOLD response for the object and color task (*bottom*) highlights a stronger and more sustained BOLD response for the object vs color task. Inset visualizes the difference between amplitude and dispersion (see Materials and Methods). (**c**) Higher BOLD dispersion (i.e., more sustained) response of V1 voxel in the object task compared to the color task. Error bars denote s.e.m; \**P* < 0.05; \*\**P* < 0.01; \*\*\**P* < 0.001.

**Figure 6.**
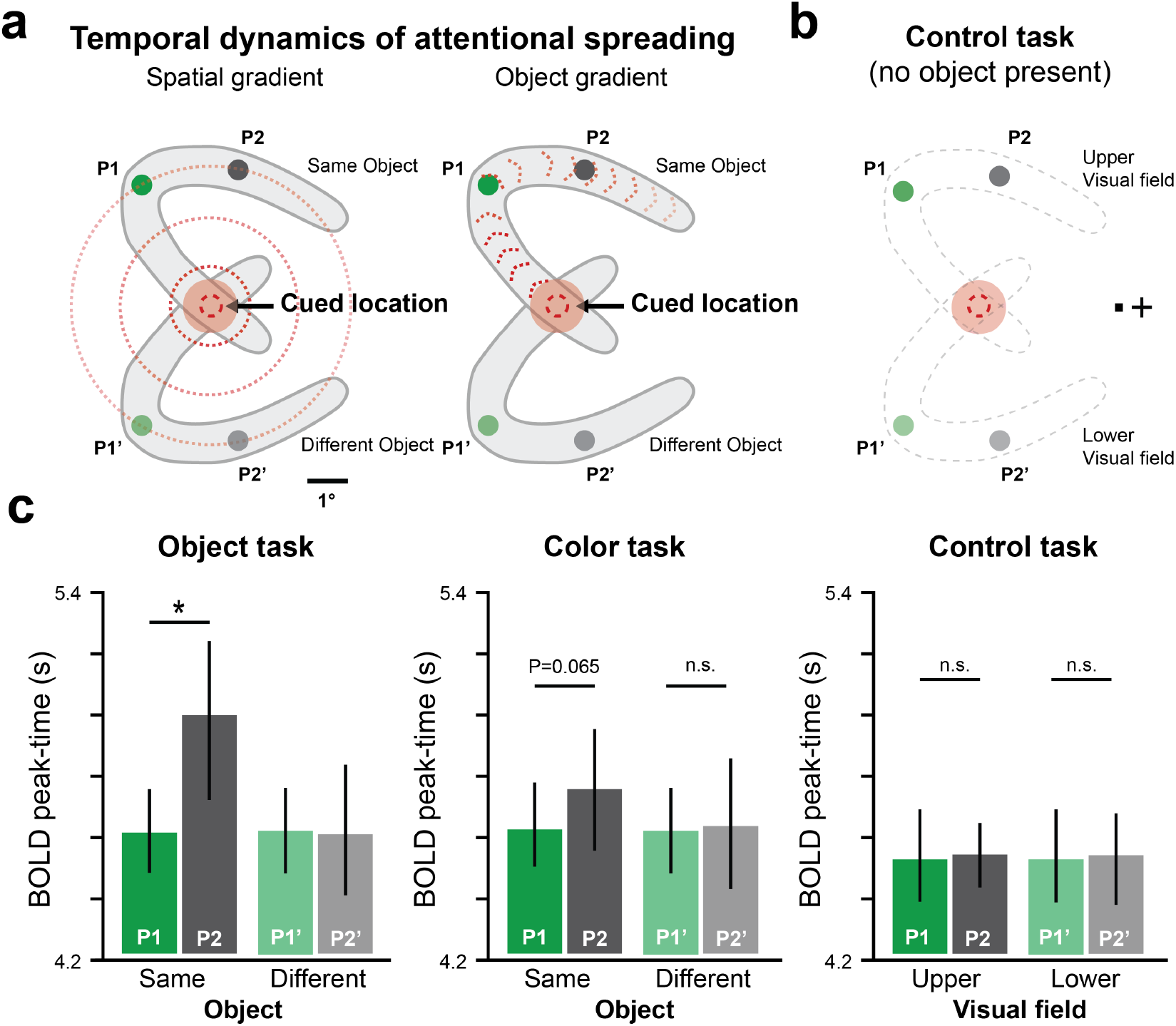
Temporal dynamics of the spread of attention follows the shape of the object. (**a**) Predictions for the spread of attention according to a spatial gradient (*left*) or along the representation of the object (*right*). If attention spreads along a spatial gradient (red dotted circles), BOLD peak-time should be identical for object locations P1 and P2. Alternatively, if object-based attention spreads over the foreground object (red dashed lines), BOLD peak-time should be delayed for object location P2 compared to P1. (**b**) Schematic of the control condition (N=7) without any objects present (grey dashed lines indicate object outline for illustration purposes only). (**c**) BOLD peak-times for the object task (*left*), color task (*middle*), and control task (*right*), separately for receptive field locations on the same object (P1, P2) and the different object (P1’, P2’), respectively. BOLD activity peaks later at object location P2 compared to P1, indicating that attention spreads across the foreground object. No differences in BOLD peak-time were observed in the control condition, and between P1’ and P2’ on the different object. Error bars denote s.e.m; \**P* < 0.05.

## Results

We quantified the spatial profile of BOLD activity in the primary visual cortex (V1) while participants viewed visual scenes composed of multiple objects (**Figure 1b**), using fMRI and population receptive field mapping. Participants were cued to direct their spatial attention to a specific location, which fell on one of four ‘horseshoe-like’ objects. Theories of object-based attention (Duncan, 1984; Chen, 2012) predict that when the intersection of the objects is cued (either on the left or right visual field), the attentional field spreads from the cued location to the foreground object, but not to the background object (**Figure 1a**). In order to test whether such attentional spreading is automatic, we further manipulated the behavioral relevance of the object. In a blocked experiment, in half of the trials participants had to decide whether two target dots appeared on the same object, thereby making the object task-relevant (“object task”). In the other half of the trials, participants had to decide whether the two presented dots were the same color, irrespective of where the dots appeared, thereby rendering the objects irrelevant (“color task”).

### Reaction times in the object task indicate serial processing

We first examined the pattern of reaction times (RTs). In accordance with previous studies (Jolicoeur et al., 1986), “same” responses, required when the second target dot was on the same (cued) object, showed faster RTs compared to “different” responses (**Figure 2b**; 728 vs 791 ms; *t*_(16)_ = −5.29, *P* = 7.3 × 10^−5^). In the color task, response were also faster if the second target fell on the foreground objects compared to the background object (690 vs 709 ms; *t*_(16)_ = −5.20, *P* = 8.7 × 10^−5^).

A possible strategy to solve the object task is to first direct attention to the cued location and then to mentally trace along the object until a second target is found (or not) (Jolicoeur et al., 1986). In contrast, the color task requires no attention to the object and mentally tracing the object is therefore unnecessary. This difference between task strategies is reflected in our data: RTs increased with the distance of two target dots measured along the object curvature. The influence of within-object distance was significant when both targets were on the same object (**Figure 2c**; average of individual correlations: *r* = 0.25, t_(16)_ = 5.91, *P* = 2.2 × 10^−5^), but not when the targets were on different objects (average of individual correlations: *r* = 0.00, t_(16)_ = 0.01, *P* = 0.99). The difference in correlation between the same vs. different object was significant (t_(16)_ = 6.14, *P* = 1.4 × 10^−5^). In contrast, RTs did not correlate with object distance in the color task (same object: t_(16)_ = 0.59, *P* = 0.57; *r* = 0.02; different object: t_(16)_ = 0.35, *P* = 0.73; *r* = 0.01). These results indicate that the appearance of the targets triggers a serial tracing process starting at the cued location and gradually spreading across the foreground object. The protracted time-course may seem surprising because the horseshoe objects were shown for a few seconds before the targets appeared, however these results are in accordance with previous findings that participants retrace objects even after repeated exposure (Wolfe et al., 2000).

### Reconstruction of Neuronal Responses Reveal Attentional Spreading

We examined neural activity in the primary visual cortex (V1) during the interval *before* the targets were shown, as we were specifically interested in the pattern of neuronal activity elicited by the spatial cue that directed attention to the left or right objects. First, we estimated the population receptive field (pRF) of every voxel in early visual cortex (see *Materials and Methods*) and calculated the BOLD activity in response to the spatial cue in comparison to baseline period without visual stimulation. Next, we projected the estimated BOLD responses from voxel space into stimulus space to examine the cueing effect on the retinotopic representation of the stimulus (Kok and De Lange, 2014; Ekman et al., 2017). Our initial reconstructions based on the independent localizer showed that the previously used reconstruction technique worked suboptimal with our curved stimuli. We therefore developed a modification of the reconstruction methods that is outlined in **Figure 3**.

Individual stimulus reconstructions were averaged across participants (N=17) for experiment 1 and revealed a clear representation of all four horseshoes (**Figure 4a**). In addition, there was enhanced V1 BOLD activity at the cued location compared to the uncued location. BOLD activity spread across the foreground object, but not across the background object. The spread of attentional over the foreground object is especially evident when we contrasted the cued (attended) vs uncued (unattended) hemifield and averaged the reconstruction over all possible stimulus configurations (**Figure 4b**; averaging across cued objects in the top/bottom and left/right hemifield). In order to statistically test the attentional spreading, we compared the neuronal response of voxels with a receptive field on the foreground object to those with a receptive field on the background object. The foreground object elicited higher activity than the background object, both during the object task (**Figure 4c**; t_(16)_ = 5.68, *P* = 3.4 × 10^−5^) and during the color task (t_(16)_ = 3.21, *P* = 5.4 × 10^−3^). The cueing effect in the object task was stronger than in the color task (**Figure 2c**; F_1,17_ = 8.19, *P* = 0.001, ANOVA (task × object)).

We found that the reconstruction also showed a representation of the objects in the unattended hemifield; however, activity differences between foreground and background objects were not significant in the unattended hemifield (F_1,17_ = 1.51, *P* = 0.24, ANOVA (task × object)), suggesting that highlighting of the foreground object depended on attention.

We further asked whether the increase in V1 activity along the foreground object was accompanied by suppression of the scene background (see **Figure 4c** for an illustration), as has been observed in area V1 of monkeys (Poort et al., 2016; Self et al., 2019). Indeed, we found significant suppression of the of the region in the center of the cued horseshoe from baseline (object task: t_(16)_ = 2.89, *P* = 0.011; color task: t(16) = 2.20, *P* = 0.043), but this effect was absent for the non-cued horseshoes (all P > .2).

### Temporal Dynamics of Object Selection Depends on Task Context

The foreground object was relevant for the object task because one of the targets appeared at the upper horseshoe at the intersection on 90% of trials and participants had to report whether the second target fell on the same object. Participants did not know when the target appeared due to the variable interval, and we assume that they maintained attention on the foreground object. Therefore, we did not expect any differences between shorter and longer delays between the onset of the cue and targets. In contrast, we predicted that if the attentional spread occurs automatically in the color task, the cueing effect on RTs would be short-lived, as the selection of one of the horseshoes had no benefit for performing the task. Therefore, we expected a difference between shorter and longer cue-target delays in the color task, with a more pronounced same-object advantage at shorter delays.

We tested this prediction by comparing RTs from shorter (i.e., shorter than the median cue-target interval duration: 1900 ms to 5150 ms; average 3525 ms) and longer (i.e., longer than median interval duration: 5155 to 7320 ms; average 6235 ms) cue-target intervals. Confirming our prediction, the same-object advantage in the color task was most pronounced at the shorter cue-target intervals (t_(16)_ = −2.86, *P* = 0.011). In the object task, the difference in RT between trials with targets on the same and different objects did not depend on the cue-target interval (t_(16)_ = −0.36, *P* = 0.723), leading to a significant task (color/object) × interval (shorter/longer) interaction (**Figure 5a**; data from experiment 1 only: F_1,17_ = 2.38, *P* = 0.13; data from experiment 1 and 2 combined: F_1,24_ = 4.32, *P* = 0.04).

In order to assess whether this behavioral effect is also reflected in the V1 neuronal dynamics, we deconvolved the BOLD activity using a finite impulse response (FIR) analysis to quantify the temporal width (i.e., dispersion) of the BOLD profile independent of its amplitude (Henson et al., 2001). Following the same logic as for the behavior, we expected a more sustained response (i.e., greater BOLD dispersion parameter) in the object task, compared to a more transient response (i.e., smaller dispersion) in the color task. Statistical comparison of the BOLD dispersion parameter confirmed this prediction (**Figure 5b/c**; t_(16)_ = 3.46, *P* = 0.0035).

### Attentional Spreading Follows the Shape of the Object

Thus far we assumed that attention spread from the cued location along the shape of the elongated foreground object (‘object gradient’). Alternatively, one could also assume that attention spread following a spatial gradient, expanding from the cued location to the visual field (Jeurissen et al., 2016). To adjudicate between these different modes of attentional spreading, we devised a second, exploratory experiment in which participants’ (N=7) neuronal responses were measured with high temporal resolution fMRI (*see Materials and Methods*). Specifically, we selected receptive fields for two locations on the object, which had the same spatial distance (~2.2 deg) but different object distance (distances measured along the medial axis of the cued horseshoe) from the cued location (points P1 and P2; **Figure 6a**). We reasoned that if attentional spreading followed a spatial gradient, BOLD peak-times for receptive field locations P1 and P2 should be indistinguishable, as both locations have the same Euclidian distance from the cued location. Conversely, if the attentional spreading followed an object gradient, BOLD peak-times for receptive field location P2 should be delayed compared to P1, as P2 was further away from the target when following the object distance. A control condition showing only target dots and omitting the objects, ensured that potential latency differences could not be simply due to inherently different BOLD peak-times at receptive field locations P1 and P2 (**Figure 6b**). Individual BOLD responses were deconvolved and fitted with a canonical hemodynamic function, and peak times were estimated independent of amplitude and baseline changes (*see Materials and Methods*).

Supporting the object gradient prediction, peak-times at location P2 were significantly delayed relative to P1 in the object task (Δt=0.43 s; t_(6)_ = 3.03, *P* = 0.0104, *BF*_*10*_ = 3.82), and showed the same – but non-significant – trend in the color task (Δt =0.22 s; t_(6)_ = 1.92, *P* = 0.0792, *BF*_*10*_ = 1.75; **Figure 5c**). No latency differences between P1 and P2 were found for the uncued object during either the object or color task (all Ps > .2).

## Discussion

The initial representation of scenes in visual cortex is heavily fragmented, as many visual features like orientation, shape and color are processed by neurons with small receptive fields distributed within and across the visual cortical hierarchy. It remains an intriguing question, how the fragmented information is integrated into a coherent representation of objects. Object-based attention theories proposed that the visual system establishes the representation of coherent objects by grouping their features incrementally with attention (Roelfsema, 2006; Roelfsema and Houtkamp, 2011). A key prediction of this account is that attention spreads from an initially selected object location to other parts of the object following low-level Gestalt cues, such as collinearity and connectedness (Wertheimer, 1923). Evidence for this mechanism comes from V1 monkey studies that observed incremental grouping of image elements of an elongated object (Pooresmaeili and Roelfsema, 2014). However, these multiunit studies can only measure from few receptive field locations simultaneously and have therefore very limited spatial resolution of the entire scene. Furthermore, neuronal correlates of the serial attentional labeling process have not yet been observed in humans.

Here we sought to test the prediction of the object-based attention account. We used fMRI to investigate V1 activity after endogenous attention was directed to a spatial location on an object in a visual scene composed of multiple horseshoe-like objects. Using population-based receptive field (prf) mapping we were able to precisely reconstruct the position and shape of the objects from V1 BOLD activity patterns, and quantify with high spatial precision the spreading of attention across the object that was cued. We observed that neuronal activity spread from the cued location across the entire foreground object in line with predictions of object-based attention account (**Figure 1a and Figure 4b)**. Additionally, in line with our predictions, a second experiment focusing on the temporal dynamics of object-based attention showed that the time-course of attentional spreading followed the shape of the cued object and not a simple Euclidian gradient.

Previous studies demonstrated that if spatial attention is summoned to a location in the visual field neuronal activity at corresponding receptive fields is enhanced (Kanwisher and Wojciulik, 2000; Pessoa et al., 2003; Sprague and Serences, 2013) and behavioral performance at the attended location is better than at unattended locations (Posner, 1990; Brefczynski and DeYoe, 1999; Posner and Gilbert, 1999). One might argue however, that most day-to-day interactions do not require attention to spatial locations, but rather to objects (e.g. looking for the coffee mug on your crowded desk). Nevertheless, the neuronal mechanisms of object-based attention remain less well understood. Our results contribute to the growing body of evidence that early visual cortex activity exceeds the representations of simple spatial locations, but also performs more complex task like object selection by means of attentional grouping.

The object task was a perceptual grouping task, because participants had to report whether two targets fell on the same or on two different objects. In accordance with previous studies, perceptual grouping in our task was associated with a gradual spread of object-attention over the relevant object so that all its features can be bound by object-based attention (Roelfsema and Houtkamp, 2011; Pooresmaeili and Roelfsema, 2014; Jeurissen et al., 2016).

In the color task, in contrast, the objects were irrelevant for the task. However, also in this task the comparison of the color of two targets on the foreground object occurred faster compared to when one of the targets fell on the background object (a ‘same-object advantage’), suggesting object-based attention had been summoned to the foreground object (Egly et al., 1994; Vecera and Farah, 1994; Müller and Kleinschmidt, 2003). Previous studies debated about the degree to which the spread of object-based attention occurs automatically (Müller and Kleinschmidt, 2003; He et al., 2004; Martínez et al., 2006; Yeari and Goldsmith, 2010; Fiser et al., 2016). This discussion has been difficult to resolve with purely behavioral methods.

A study by Müller & Kleinschmidt (2003) reported that activity along other object locations was only increased if targets did not appear at the expected object location. The authors argued that their findings are in line with theories that cast attentional spreading not as automatic, but rather as an “object-based search strategy” (Moore et al., 1998; Shomstein and Yantis, 2002); here activity spreads along the object, but only if the relevant information is not found at the attended location. In disagreement with these results, a recent study in V1 monkeys demonstrated that neuronal activity automatically spreads from attended image element to other image element that are grouped by Gestalt criteria (Wannig et al., 2011). Our present results support the automatic spreading account by demonstrating the spreading of extra BOLD activity over the entire foreground object also in the color task. Notably however, in the color task the behavioral “same-object advantage” was more transient and the same was true for the increased BOLD activity along the cued object, which was of shorter duration than in the task in which the objects were task-relevant.

In sum, the current study provides evidence for automatic attentional spreading from a spatial location to an underlying object. Attentional selection can be reconstructed with high spatial fidelity and reveals an interaction of endogenous spatial attention and exogenous object selection that is independent of task context. The current study thereby provides empirical support for theories that cast attentional object-based spreading as an automatic process.

## Author contributions

M.E., P.R.R. and F.P.d.L., conceived and designed the experiments. M.E. collected the data and conducted the data analyses. M.E., P.R.R. and F.P.d.L. wrote the manuscript.

## Acknowledgements

This study was supported by the Netherlands Organization for Scientific Research (Veni Grant No. 016.Veni.195.435), awarded to M.E., the James S. McDonnell Foundation (JSMF Scholar Award for Understanding Human Cognition), the European Union Horizon 2020 Program (ERC Starting Grant 678286, “Contextvision”), awarded to F.P.d.L., the European Union’s Horizon 2020 and FP7 Research and Innovation Program (grant agreement ‘‘Human Brain Project SGA2’’, ERC grant agreement 339490 ‘‘Cortic_al_gorithms’’ awarded to P.R.R., and the Friends Foundation of the Netherlands Institute for Neuroscience. We thank Alya Vlassova and Micha Heilbron for comments on an earlier version of the manuscript.

